# HIV-1 infection of genetically engineered iPSC-derived central nervous system-engrafted microglia in a humanized mouse model

**DOI:** 10.1101/2023.04.26.538461

**Authors:** Alice K. Min, Behnam Javidfar, Roy Missall, Donald Doanman, Madel Durens, Samantha St Vil, Zahra Masih, Mara Graziani, Annika Mordelt, Samuele Marro, Lotje de Witte, Benjamin K. Chen, Talia H. Swartz, Schahram Akbarian

## Abstract

The central nervous system (CNS) is a major human immunodeficiency virus type 1 reservoir. Microglia are the primary target cell of HIV-1 infection in the CNS. Current models have not allowed the precise molecular pathways of acute and chronic CNS microglial infection to be tested with in vivo genetic methods. Here, we describe a novel humanized mouse model utilizing human-induced pluripotent stem cell-derived microglia to xenograft into murine hosts. These mice are additionally engrafted with human peripheral blood mononuclear cells that served as a medium to establish a peripheral infection that then spread to the CNS microglia xenograft, modeling a trans-blood-brain barrier route of acute CNS HIV-1 infection with human target cells. The approach is compatible with iPSC genetic engineering, including inserting targeted transgenic reporter cassettes to track the xenografted human cells, enabling the testing of novel treatment and viral tracking strategies in a comparatively simple and cost-effective way *vivo* model for neuroHIV.

**Importance:** Our mouse model is a powerful tool for investigating the genetic mechanisms governing CNS HIV-1 infection and latency in the CNS at a single-cell level. A major advantage of our model is that it uses iPSC-derived microglia, which enables human genetics, including gene function and therapeutic gene manipulation, to be explored *in vivo*, which is more challenging to study with current hematopoietic stem cell-based models for neuroHIV. Our transgenic tracing of xenografted human cells will provide a quantitative medium to develop new molecular and epigenetic strategies for reducing the HIV-1 latent reservoir and to test the impact of therapeutic inflammation-targeting drug interventions on CNS HIV-1 latency.

## Introduction

The pandemic of human immunodeficiency virus type 1 (HIV-1) causes a chronic infection without a known universal cure (1). While there have been three reported cases of HIV-1 remission and cure after bone marrow and cord blood transplantations, these procedures are extremely high risk and have been developed for rare cases of patients with life-threatening, co-morbid leukemia (2). A significant challenge to finding a sterilizing cure has been the persistence of latently infected cellular reservoirs that resume replication when antiretroviral therapy (ART) is withdrawn (3–8). Furthermore, while ART can suppress viremia to undetectable levels, people with HIV-1 (PWH) experience more premature aging and inflammation-associated pathophysiology than uninfected people (9–13). The central nervous system (CNS) carries a heavy disease burden with HIV-1-associated neurocognitive disorder, affecting 20-50% of PWH (14–16).

HIV-1 enters the brain during the first initial weeks of acute infection (17–20) via transmigration of HIV-1 infected cells across the blood-brain barrier (BBB) (21–23). In the periphery, HIV-1 targets the CD4^+^ T cells and monocytes that then disseminate infection to other tissues and organs (21–26). In the CNS, these peripherally infected cells target primarily microglia comprising about 5-10% of adult brain cells (27). Our recent studies on the human postmortem brain corroborate that HIV-1 infection occurs predominantly in microglia, and provirus integration is linked to inflammation-associated reprogramming of microglial transcriptomes and 3D genomes (28). Understanding the molecular mechanisms underlying the establishment and maintenance of actively and latently HIV-1 infected microglia in the CNS will help investigate ways to target and reduce CNS reservoirs and develop therapeutic interventions for microglia-associated neuroinflammation.

Genetic animal models that capture the salient features of CNS HIV-1 infection in humans are greatly needed to interrogate the precise mechanisms governing the HIV-1 life cycle in the host. These can enable the study of initial infection of the peripheral immune system, viral replication, dissemination throughout all organ systems, including the CNS microglia and other myeloid compartments, and provirus integration into the host cell genome to maintain chronic HIV-1 infection. To this end, elegant non-human primate models using various strains of Simian Immunodeficiency Virus (SIV) have long been established in the field (29, 30). However, these models are still inherently limited, given that HIV-1 is a human-specific retrovirus (31).

Although HIV-1 does not replicate in mouse immune cells, the repertoire for modeling neuroHIV has been expanded by employing chimeric virus models to infect mice. The EcoHIV model utilizes a chimeric murine-tropic virus that has been genetically modified to carry a Murine Leukemia Virus envelope coding region and hence can infect and simulate HIV-1 disease-like phenotype in mice (32). To study native HIV-1 in small animal models, investigators have employed human immune cell xenografted immunodeficient mouse models. While initially, studies focused on blood and peripheral systems (33, 34), a recent advance has enabled CNS engraftment of human cells by using human fetal hematopoietic tissue or cord blood-derived CD34^+^ hematopoietic stem cells (HSC) into immunocompromised NOG mice expressing a human Interleukin(IL)-34 transgene. These mice show colonization of multiple lymphoid and myeloid compartments of host mice, including differentiation of microglia-like brain cells susceptible to HIV-1 infection (35, 36). Recently, a variation of this model where the mice are xenografted instead with human fetal liver tissue, fetal thymic tissue, and liver-derived CD34^+^ hematopoietic stem and progenitor cells (HSPC) was reported to undergo human microglia reconstitution in CNS that are susceptible to HIV-1 infection (37). Other models rely on the direct engraftment of HIV-1-infected human myeloid cells into the brain of immunocompromised mice (38).

These humanized mouse models have been extremely valuable for expanding knowledge on molecular and cellular signatures of HIV-1 infected brains. However, currently, existing humanized mouse models also pose some limitations. These include the requirement of human fetal tissue or umbilical cord blood that are limited in supply and not readily amplified to conduct extensive tests with the same genotype (35, 37, 39) and exhibit a lower rate of xenograft reconstitution following irradiation or chemotherapy (35–37) as compared to iPS-derived cells. Additionally, some of these models feature a non-physiological intracranial route of HIV-1 inoculation into the brain (35, 38, 40). Here, we present a novel humanized mouse model for HIV-1 infection that implements genetically engineered human induced pluripotent stem cells (iPSCs) as a near-unlimited resource to reconstitute neonatal mice brains with human microglia. The unrivaled versatility of iPSCs enables genetic material to be clonally engineered and provides an invaluable toolbox for generating reporter cell lines. In our model, CNS-engrafted mice are peripherally engrafted with human peripheral blood mononuclear cells (PBMC) in early adulthood to initiate infections from peripheral vasculature. Infected PBMCs then travel to the CNS to infect the xenografted microglia (xenoMG). Mice dually engrafted with xenoMGs and PBMCs are infected with M-tropic HIV-1. This models a physiological route of acute CNS HIV-1 infection through virus or infected cells crossing the BBB.

## Results

### Differentiation of microglia from genetically modified human iPSC in vitro

Studying CNS HIV-1 infection has been challenging due to the limited accessibility of human samples and difficulty in tracing persistently infected cellular reservoirs, even with suppressive ART. Here we present a novel human iPSC-based mouse model that utilizes Cre-recombinase-dependent lineage tracing to mark cells that have been HIV-1 infected irreversibly. We genetically engineered the well-characterized human induced pluripotent stem cell (iPSC) line WTC11 derived from an apparently normal individual(41), inserting a Cre-recombinase dependent, CAG promoter-driven dsRed-to-eGFP switch cassette into the AAVS1 locus within the *PPP1R12C* gene (MSE2104, Supplemental Fig 1A). The AAVS1 locus is an exemplary site for genetically engineering iPSC because it permits robust, stable, and reproducible expression of transgenes along multiple lineages maintained long-term in culture (42). It also overcomes the pleiotropic position effects, and the inserted transgene is not silenced during meso- and ectoderm differentiation. The genetic switch is encoded by flanking *dsRed* stop with *loxP* cre-recombinase target sites upstream of an *egfp* coding sequence, enabling cre-dependent switching of dsRed to eGFP (Supplemental Fig 1A, B).

The iPSCs were differentiated into hematopoietic progenitor cells (HPCs) using a commercially available STEMdiff Hematopoietic kit as per published protocols (Figure 1A) (43–47). The development of HPCs from iPSCs requires a combination of growth factors, cytokines, and small molecules, which include stem cell factor (SCF), Flt3 Ligand (Flt3L), Thrombopoietin (TPO), IL-3, IL-6, and Granulocyte Macrophage Colony-Stimulating Factor (GM-CSF) (48, 49).

**Figure 1:**
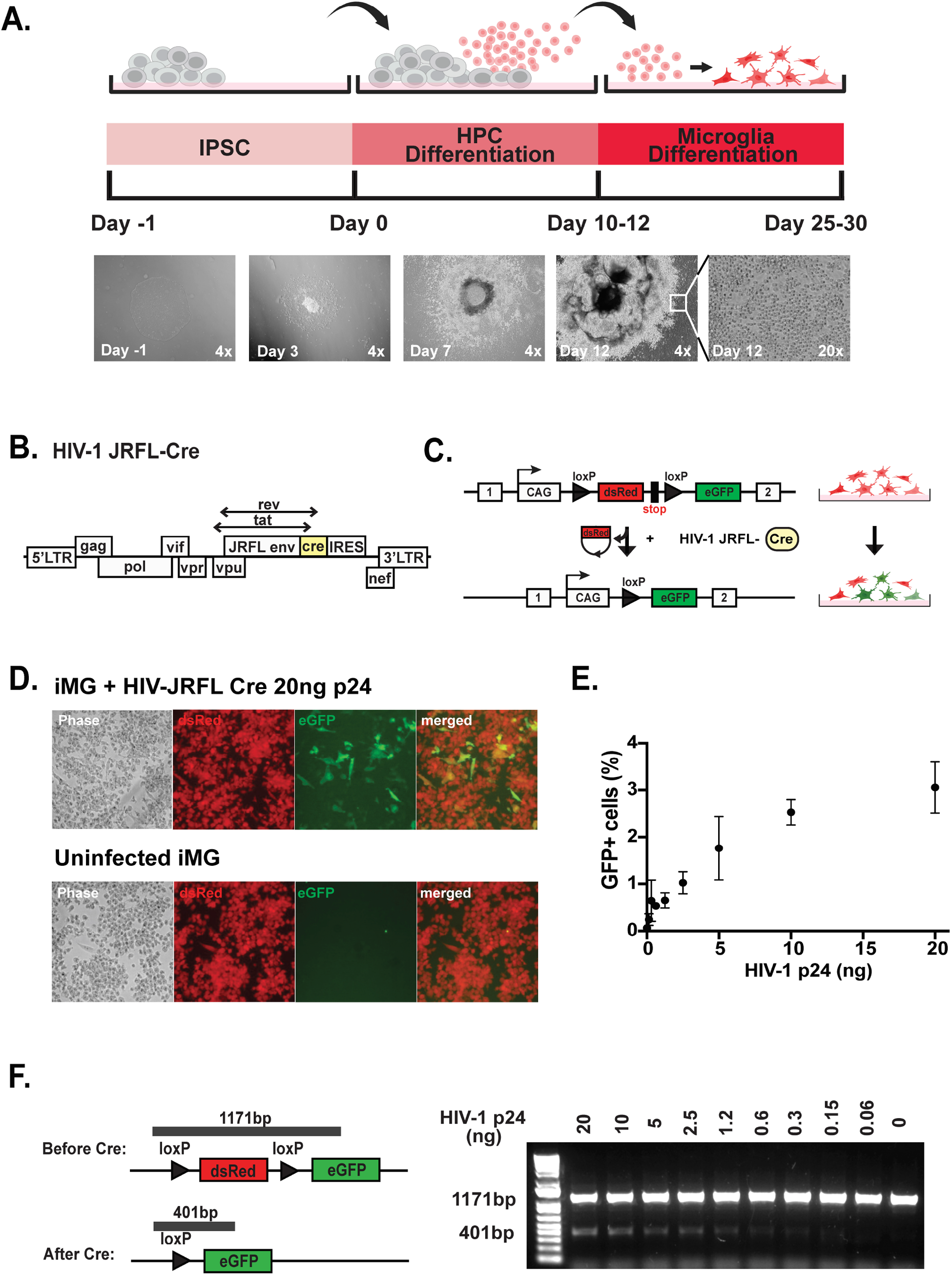
Overview of in vitro experiments: (**A**) (top) Timeline of iPSC-HPC-iMG differentiation with (bottom) corresponding, stage-specific morphological appearances of cell cultures at low power magnification (4x – 20x, as indicated) of (top) conditional reporter transgene insertion at chromosome 19 AAVS1 (Adeno-Associated Virus Integration Site 1) targeting locus. (bottom). (**B**,**C**) Schematic of (B) HIV-1 JFRL-Cre genome including Cre coding cassette followed by IRES internal ribosomal entry sequence to drive *nef* expression for conditional recombination of the dsRed-GFP reporter and (C) dsRed to eGFP switch in HIV-Cre infected cells. (**D**) Conditional transgene expression: with (top) HIV-1 JFRL-Cre infected and bottom (uninfected) MSE2104 iPSC-derived microglia (iMG). Notice robust ubiquitous dsRed and, in a subset of cells, GFP expression. Cells were harvested 48h post-infection with 20ng p24 HIV-1 JRFL-Cre virus. (**E**) Approximately 3% of GFP+ MSE2104 iMG with 20ng p24 HIV-1 JRFL Cre. (**F**) Recombination-sensitive 401bp PCR product extending into 5’ of GFP cassette, as indicated. Notice the dosage-sensitivity of recombination to virus levels (p24 range 0-20ng) in cell culture, as indicated.

Differentiating iPSC into HPC took ten days, after which cells were harvested between days 10-12 for downstream experiments. We validated our HPC differentiation by measuring established HPC-specific CD34, CD43, and CD45 cell surface marker expression on flow cytometry. As expected, more than 90% of HPCs expressed CD34 and CD43, and approximately 50-60% of HPCs expressed CD45 (Supplemental Fig 2A) (43–45, 50).

To generate homeostatic microglia (iPSC-derived MG, or iMG) in vitro, HPCs were treated with cytokines IL-34, Transforming Growth Factor beta 1 (TGFβ-1), and Macrophage Colony-Stimulating Factor (M-CSF), promoting the development of (Figure 1A, Supplemental Fig. 2B) (43–47, 50). Differentiation of HPC to iMG was followed for 20-25 days, and phenotypes were validated by measuring the surface expression of common myeloid marker CD45 and microglial markers, Iba1 and P2YR12. Macrophage-specific CD206 and monocyte-specific CD14 cell surface markers were examined to distinguish our iMGs from macrophages and monocytes. More than 90% of our iMGs expressed CD45 and CD11b, and approximately 80% expressed P2YR12 (Supplemental Fig 2B). Furthermore, most of our WTC11 iPSC-derived iMGs stained positive for P2YR12, Iba1, and CD45 on immunofluorescence (Supplemental Fig 2C).

### Cre-activated GFP Reporter iPSC-derived microglia is infected by M-tropic HIV-1 clone in vitro

To determine the ability of in vitro differentiated iPSC-derived cells to support HIV-1 infection, we constructed an HIV-1 clone that carries the M-tropic JRFL envelope to target all myeloid lineage cells, including microglia (Figure 1B) (51–53). This virus also expressed Cre instead of the viral Nef, an early viral gene expression indicator. Cre-recombinase-dependent cellular changes occurred only after HIV-1 viral integration (54). Nef expression was restored by inserting an internal ribosome entry site (IRES) upstream of the Nef open reading frame, as previously described (54). The iMGs were infected with 20ng p24 of HIV-1 JRFL-Cre, as previously described (53–56). After 48 hours, HIV-1 JRFL-Cre infected cells started to express eGFP (Figure 1C, D). On average, we saw approximately 3% of infected cells (Figure 1E). Hence, many cells continued to express dsRed, and some even appeared to co-express dsRed and eGFP. This dsRed, and eGFP co-expression by infected iMGs is likely because eGFP is expressed early upon provirus integration and the potentially long maturation time and half-life of dsRed (57, 58). The Cre recombinase-mediated deletion of dsRed in infected cells was confirmed by DNA PCR (Figure 1F). Our fluorescent reporter IPSC line can be successfully used to generate human microglia (iMGs) and could serve as a tool for studying HIV-1 infection at a single-cell level *in vitro* and, as described below, *in vivo*.

### A novel humanized mouse dually xenografted with human iPSC-derived microglia and PBMC

Next, to generate mice with iPSC-derived microglia, we engrafted iPS-derived precursor cells into immunocompromised mice harboring the human mCSF (*CSF1*) knock-in allele in a *Rag2* and *Il2rg* knockout background. These mice express at least one allele of the human *CSF1* gene that critically supports the development of human iPSC-derived microglia in mice brain (43–47, 59). Mice lacking *Rag2* and *Il2r*g genes have no native T cells, B cells, and NK cells; hence, they are ideal hosts for xenografted human cells. Genetically-modified fluorescent reporter iPSC described above were differentiated into HPCs and injected intracranially into newborn pups on post-natal days 0-2 (Figure 2A). At three months, we observed widespread distribution of dsRed+ xenografted cells in the forebrain cerebral cortex of the mice (Figure 2B). To confirm that the engrafted dsRed+ cells are human and differentiated into microglia *in vivo*, we labeled sections from mouse brains at post-engraftment week 8 for expression of the Ku80 human-specific nuclear antigen (44) and microglia-specific cell markers. Indeed, the dsRed+ cells colocalized with Ku80 and microglia-specific ionized calcium-binding adapter molecule 1 (Iba1) and purinergic receptor P2RY12 cell markers (Figure 2C), confirming that the engrafted hIPSC-HPC had differentiated into microglia/(brain) tissue-resident macrophages.

**Figure 2:**
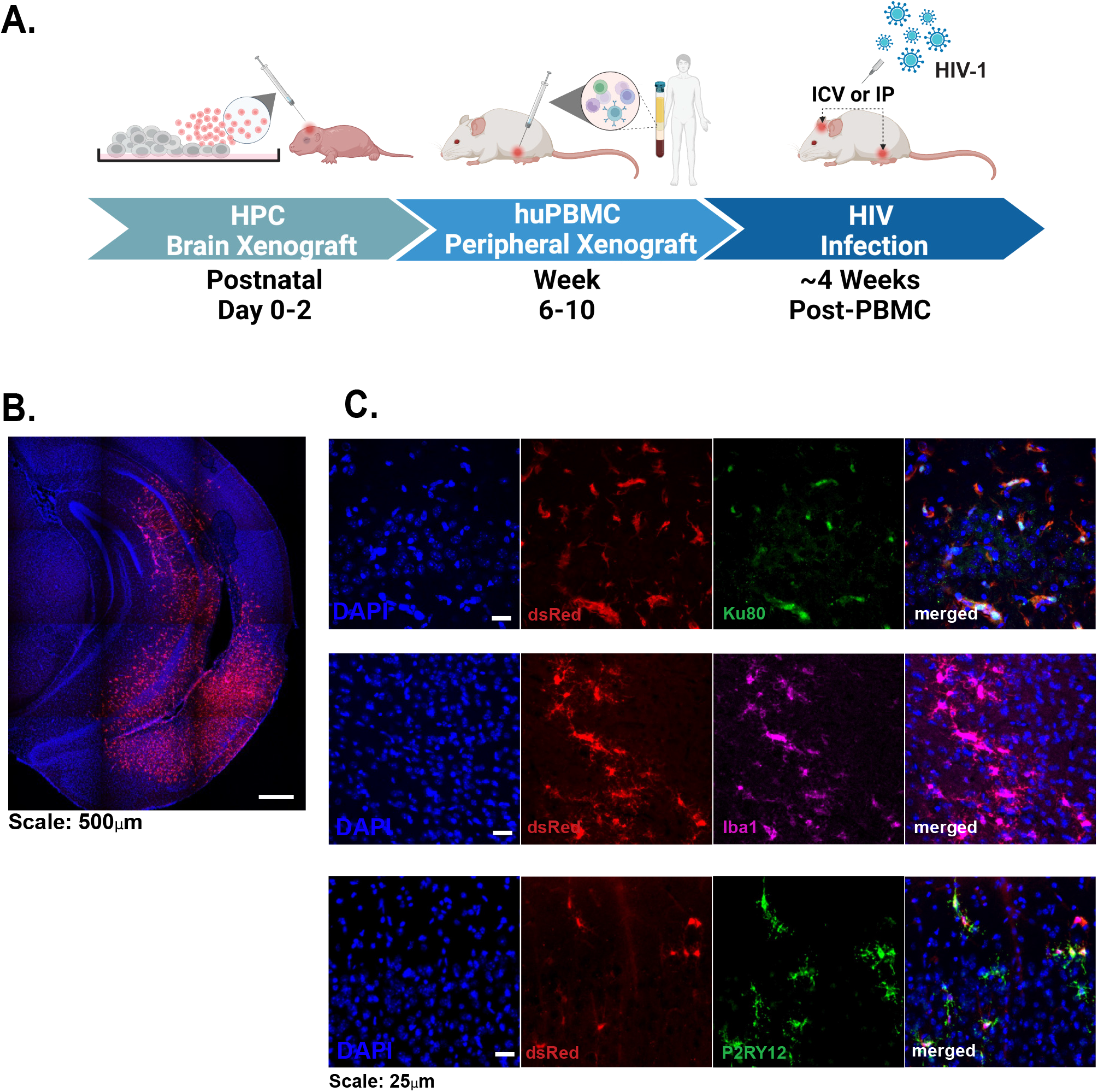
Humanized mouse model with central and peripheral xenograft. **(A)** Timeline and overview, a central hiPSC-HPC xenograft into the neonatal brain is followed by peripheral hu-PMBC administration (i.p.) in young host animals, followed four weeks later by central or peripheral infection with HIV. (**B**) 4x magnification of whole hemisphere view of DAPI-counterstained (blue) coronal brain section from adult animal neonatally engrafted with dsRed+ hIPSC-HPC microglial progenitors. Notice large numbers of dsRed-expressing cells at particularly high densities in the medial temporal lobe, including the hippocampus. (**C**) 40x magnification from coronal sections at upper layers I and/or II of the adult cerebral cortex of neonatally engrafted animals confirms numerous dsRed-expressing human cells and myeloid markers Iba1 and P2RY12 purinergic receptors. DAPI, counterstain. Scale bars (A) 500, (B) 25 μm.

In our mouse model, we aimed to infect mice via peripheral immune cells that cross the blood-brain barrier (BBB) rather than through a non-physiological direct injection of virus intracranially (Figure 2A). To accomplish this, in addition to the neonatal brain engraftment with human iPSC-derived microglia (xenoMG), we peripherally engrafted our mice in early adulthood (postnatal week 6-10) with human peripheral blood mononuclear cells (huPBMC). This dual engraftment allows us to model the physiological route of acute HIV-1 infection of the CNS in humans, with the peripheral blood CD4+ T-cells and monocytes as initial primary targets. Each mouse was intraperitoneally injected with phytohemagglutinin-L (PHA-L) and Interleukin (IL)-2 activated 10^7^ huPBMCs (see methods). Post-injection, huPBMC engraftment efficiency was monitored by cheek blood flow cytometry for expression of human leukocyte-specific cell surface markers: leukocyte common antigen CD45 and T cell markers CD3, CD4, and CD8 (Supplemental Fig 3). We expected a comparatively lower abundance of human monocytes (data not shown) in the mice blood relative to human CD45^+^ T cells based on the low expression (∼10-15%) of monocyte cell markers CD14 and CD16 in pre-injected huPBMC (Supplemental Fig 4). In this model, mice are expected to reach a minimum of 20% human immune cell engraftment within the pool of circulating leukocytes in the mice blood approximately four weeks post-huPBMC injection.

We next tested the ability of these dually engrafted mice to be infected with our HIV-1 JRFL Cre clone. We investigated two different routes of infection: intracerebroventricular (ICV, n=4) and intraperitoneal (IP, n=5) (Figure 2A, Figure 3A). We monitored peripheral IP HIV-1 infection by weekly monitoring of HIV-1 viral load via RT-qPCR in mice engrafted with huPMBCs (Supplemental Fig 5A) and control mice with no huPBMCs. As expected, we observed that huPBMC-engrafted mice have HIV-1 copy numbers at least 2-3 orders of magnitude above the detectability threshold compared to controls not engrafted with PBMCs (Figure 3A). Specifically, the HIV-1 copy number in our huPMBC engrafted mice ranged from ∼10-1000 viral RNA transcript copies/ml blood, a number that is comparable to early stages of HIV-1 infection in human subjects whose average viral load is 30,000 to 50,000 copies/mL before initiating effective antiretroviral therapies (ART) (60–62). Likewise, ICV-infected mice receiving PBMC transplants showed higher HIV-1 viral load than the no PBMC control cohort (Fig. 3A). HIV-1 viremia peaked three to four weeks post-infection (Supplemental Figure 5), at which point the brains were harvested and processed for immunohistochemistry.

**Figure 3:**
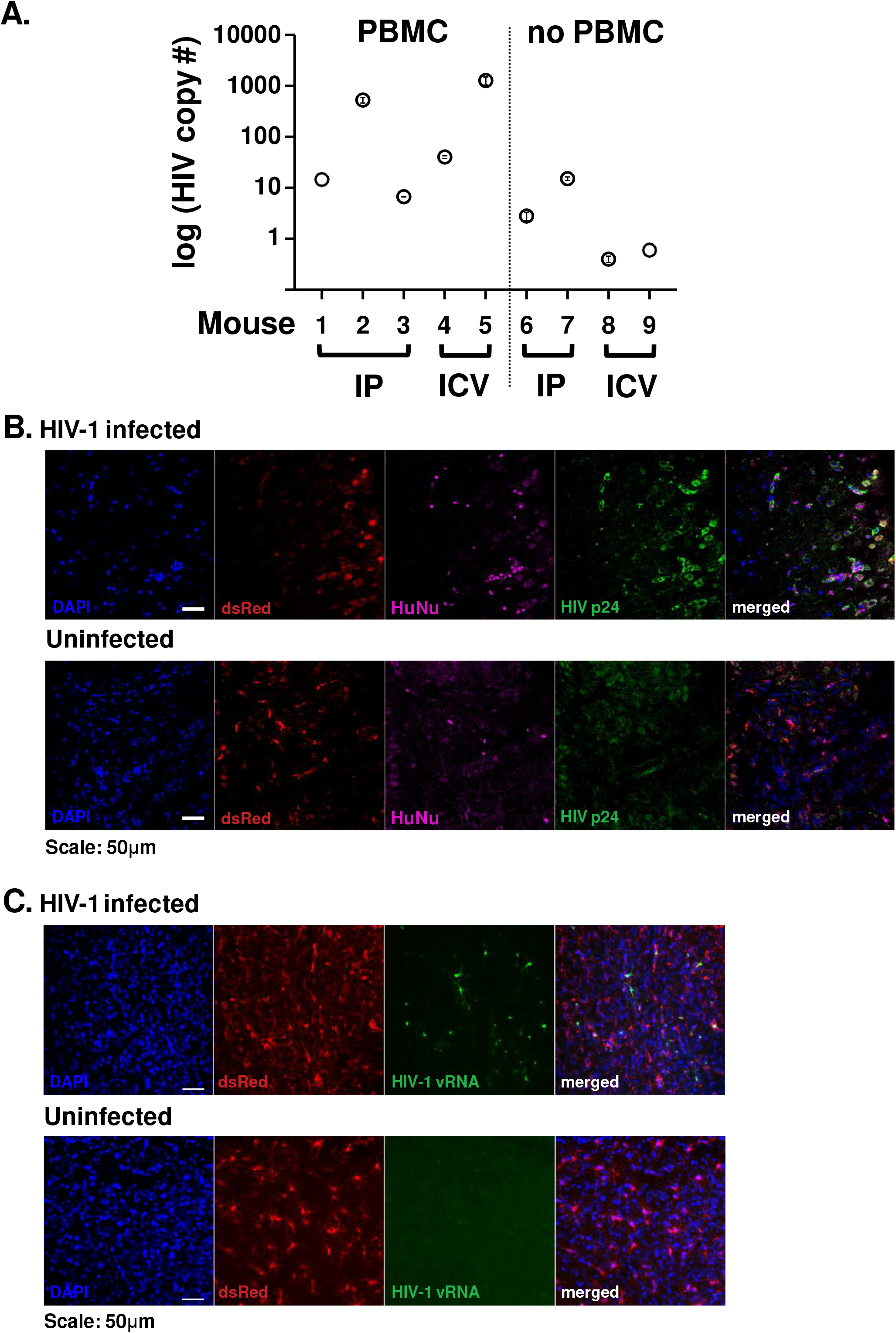
Neonatally engrafted hIPSC-HPC microglia become HIV infected after peripheral infection. (**A**) (y-axis) Quantitative PCR-based blood HIV RNA copy numbers per ml cheek blood, (x-axis) in individual mice no. 1-9 neonatally engrafted with hIPSC-HPC, including with (no. 1-5) and without (no. 6-9) additional huPMBC transplant as an adult. All Animals had received a single dose of HIV (250ng p24 antigen) i.p., four weeks before the first blood draw. N= 2-3 blood draws/animal, data shown as mean ± S.E.M. Notice robust titers are limited to animals that received hu-PMBC. (**B,C**) Representative sections from the adult rostromedial cerebral cortex of neonatally hiPSC-HPC and adult hu-PBMC engrafted mice, (top) HIV infected (bottom) uninfected animals. Sections were counterstained nucleophilic dye DAPI (blue) and show human iMG expressing dsRed reporter transgene, with additional co-staining by (**B**) immunohistochemistry with HuNu human-specific nuclear antigen to mark human cells and HIVp24 antigen (green) and (**C**) HIV gag-pol RNA FISH (green) with the latter showing above background signal in the infected mouse. Scale bars, 50 μm.

We focused our histological analysis initially on N=3 dually engrafted mice subjected to HIV-1 infection via IP route. The brain tissue sections of these mice revealed human nuclear antigen-positive (HuNu) human nuclei surrounded by a halo of HIV-1 p24 antigen in the cytoplasm, confirming successful infection (Figure 3B). Remarkably, a subset of HIV p24^+^ cells presented as large multinucleated HuNu^+^ cells, reminiscent of the well-described multinucleated microglial nodules of HIV-1 infected cells in the human brain, particularly in cases with severe neuroinflammation and encephalitis (63, 64). In addition, HIV-1-infected mice brains had xenoMGs with a more amoeboid shape where the soma became more enlarged, and the cells had no or fewer processes that appeared shorter and wider, reminiscent of similar findings in clinical specimens (63, 65, 66). In contrast, the xenoMGs in uninfected mice brain tissue resembled the typical homeostatic microglia morphology with smaller soma, more and longer processes with more ramifications (Figure 3B). Finally, for additional confirmation of successful xenoMG infection with our strain of HIV-1, we performed RNAScope fluorescence in situ hybridization (FISH using HIV-1 specific RNAscope probes). There was robust expression of HIV-1 viral RNA in the xenoMGs of HIV-1 infected mice. In contrast, no signal above the background was detectable in the brains of uninfected mice (Figure 3C).

## Discussion

Here, we present a novel humanized mouse model where mice are dually engrafted with human iPSC-derived, genetically-engineered microglia centrally and huPBMC peripherally. While the brains of these mice can be infected with HIV-1 via ICV or IP injection, the peripheral route offers the key advantage of modeling the typical route of CNS infection in human cases. A major advantage of our model is the use of highly versatile iPSC that could be used as a tool for further genetic engineering, including the transgenic reporter system that allows us to track the transplanted cells long-term, even four months after transplantation (the terminal time-point in our study) into the neonatal murine CNS.

Utilizing iPSCs to differentiate human microglia and subsequent xenograft into the mouse brain offers an abundant and renewable source of microglia. It also permits genetic modification before transplantation and *in vivo* experimentation, paving the way for a more sophisticated understanding of HIV-1 pathogenesis. The use of genetically modified stem cells has proven immensely valuable in HIV-1 research, facilitating the assessment of cellular susceptibility and resistance to infection by changing viral docking sites on the cell surface (67) and enabling the investigation of cell type-specific transcriptional responses to infection and antiretroviral treatment in a tightly controlled, isogenic environment (68, 69). Despite the critical importance of these molecular and mechanistic studies, they have primarily been conducted *in vitro* or *ex vivo*, which can limit their clinical relevance. Our novel humanized mouse model presents a promising alternative for preclinical and translational work *in vivo*, providing a platform for assessing genetically engineered human iPSC-derived cells in the brain, thereby unlocking new frontiers for developing more effective HIV-1 therapies.

To develop novel therapies for treating and potentially eradicating HIV-1, we propose utilizing genetically edited iPSCs targeting promising genes. For instance, the CCR5 Δ32 mutation has exhibited substantial promise as a curative target against HIV-1, as demonstrated by multiple clinical studies indicating its high resistance to the virus (70–74). Similarly, SAMHD1, a host restriction factor that impedes HIV-1 reverse transcription in myeloid cells, presents a potential therapeutic target for combating HIV-1 (75–79). Thus, our genetically engineered iPSC-based mouse model significantly broadens the repertoire of humanized mouse models for CNS HIV-1 infection, augmenting our capacity to explore innovative approaches to combat this debilitating disease.

Several other humanized mouse models have been developed to study CNS HIV-1 infection (35-37, 80-83). The hIL34-NOG model involves intrahepatic transplantation of umbilical cord blood-derived CD34^+^ HSPCs that differentiate into human microglia in mice brains over 6-8 months before infection with HIV-1 (35, 36, 81). The hu-BLT-hIL34-NOG is a modification of the hIL34-NOG model that involves transplantation of human fetal liver, thymic tissue, and fetal liver-derived CD34^+^ HSPCs. This model reconstitutes the human immune system in the mouse by 16 weeks of age, and the mouse is susceptible to HIV-1 infection (31, 37, 40). However, these models are significantly limited by their restricted options for genetic manipulation. Our iPSC-based humanized mouse model allows clonal expansion of genetically modified iPSCs with uniform penetrance of the altered gene. This holds tremendous potential for generating knock-out, knock-in, or conditional transgenes to definitively examine the role of genes within an in vivo context.

In addition, both models require several months for complete reconstitution of the human hematolymphoid system before effective HIV-1 infection (84). Our iPSC-based humanized mouse model offers a viable alternative for achieving CNS HIV-1 seeding within a relatively shorter time. Our iPSC-based humanized mouse model utilizes huPBMCs that engraft rapidly within four weeks and efficiently in immunocompromised mice, thus shortening the time from engraftment to infection analysis compared to other humanized mouse models. Moreover, this approach also allows simulation of physiological peripheral HIV-1 infection during acute infection in humans, where HIV-1 infects immune cells such as CD4^+^ T cells or monocytes that cross the blood-brain barrier into the brain, subsequently targeting microglia to disseminate HIV-1 infection in the CNS (85–90). Hence, our humanized mouse model dually engrafted with xenoMG and huPBMC provides a faster alternative for achieving CNS HIV-1 seeding.

On the other hand, immunocompromised mice engrafted with huPBMC, as shown here, generate a T cell dominant model. While this model supports robust and rapid acute HIV-1 infection kinetics, it lacks specialized immune cell subsets found in fetal or cord blood-derived immune cells, producing a less diverse immune system model. Umbilical cord blood and fetal tissues, rich in hematopoietic stem cells and have a high capacity for tissue regeneration, allow for a more complete reconstitution of the human immune system and create a more robust humanized mouse model for studying diseases. Our iPSC-derived xenoMG system, which relies on T cell-dependent infiltration of HIV-1 into the CNS, provides a platform and starting point for investigating HIV-1 infection. Future alternative approaches, such as transplantation of cord blood-derived HSC instead of huPBMC, will further diversify and strengthen the iPSC-based humanized mouse model for studying CNS HIV-1 infection.

Despite confirming HIV-1 expression through various methods, our Cre-activated GFP reporter transgene expression has not been readily detected in the humanized mouse brain at 30 days post-infection (data not shown). Possible explanations may include inefficient maturation of the GFP fluorophore and or weaker transgene expression in HIV-infected microglial cells. In other studies of acute infection in humanized mice, the Cre-activated switch effectively marks HIV DNA-positive cells for several weeks in T cells (Satija et al. in preparation). However, to further develop the use of the *in vivo* Cre-activated switch for xenoMG, additional studies will be required to follow the maintenance of Cre-sensitive sequences over time. The presence of the reporter transgene in the virus may have resulted in a metabolic burden during ongoing replication, potentially impacting the viral fitness and resulting in the deletion of the transgene. While our results highlight challenges associated with using a Cre-activated reporter to track HIV-1 infection *in vivo*, further studies are needed to fully explore this approach’s potential and feasibility.

A Cre-activated reporter-dependent humanized mouse model presents a powerful tool for investigating the pathogenesis of both active and latent HIV-1 infection (Satija et al. in preparation). The Cre-loxP system allows for the tracking of HIV-1 infected cells *in vivo*, enabling the characterization of viral gene expression and the identification of cellular reservoirs of latent HIV-1. In any case, transgenic reporter systems, including the one presented here, can be utilized to track human cells including xenoMG in the animal host, which also will be convenient for single-cell RNA sequencing analysis, chromatin analysis, epigenetic and molecular analysis of HIV pathogenesis, and the testing of therapeutic drug candidates against HIV. Understanding the mechanisms of HIV-1 latency is crucial for developing strategies to eliminate latent viral reservoirs, which are a major barrier to an HIV cure. Thus, utilizing humanized mouse models with genetically engineered hIPSCs to study HIV-1 latency could provide invaluable insights into the pathogenesis of HIV-1 and the development of effective treatments for HIV-1 infection.

## Acknowledgments

The authors thank Joe Castellano and Brittany Hemmer for their valuable input and help on humanized mouse model development and Susan Morgello for critical input about HIV-1 neuropathology. We also thank Alice Buonfiglioli and Daniele Mattei for their help, discussion, and advice on microglia work.

## Funding

Supported by NIDA R01 DA054526 and DP1056018 and NIAID R01 AI162223.

## Data availability/accessibility

All protocols and materials will be provided by the Authors upon request.

## Conflicts of Interest

The Authors do not declare a conflict.

## SUPPLEMENTARY FIGURE LEGENDS

**Supplementary Figure 1**: **MSE2104 iPSC line (genetically engineered WTC11 cells) with conditional reporter transgene**. (**A**) overview on Cre-sensitive reporter system, showing (**B**) by FACS analysis and (**C**) by phase contrast microscopy, GFP+ cells after transfection with Cre plasmid in MSE2104 but not in untreated control cultures.)

**Supplementary Figure 2**: **MSE2104 hiPSC differentiate in vitro into hematopoietic progenitor cells (HPC) and microglia (iMG)**. (**A, B**) Flow cytometry-based proportional representation of (**A**) CD34, CD43, and CD45 cell surface markers in hIPSC-HPC and (**B**) CD45, CD11b, CD14, CD206, and P2YR12 cell surface markers in hiPSC-HPC-iMG differentiated MSE2104 cells. (**C**) Immunocytochemical staining of hIPSC-HPC-iMG cultures stained with CD45 and myeloid/microglia-specific markers Iba1 and P2RY12.

**Supplementary Figure 3**: **Blood T-lymphocyte quality control check**. (top) Mouse peripheral cheek blood sampled four weeks after receiving huPMBC xenograft, notice a sizeable portion of CD4 and CD8 lymphocyte fraction. (bottom) Representative cell surface marker expression profile of CD3 gated CD4 and CD8 lymphocyte population from activated (pre-injection) huPBMC sample.

**Supplementary Figure 4: Blood monocyte/macrophage quality control check**. The activated (pre-injection) huPBMC sample shows a proportional representation of CD14 and CD16 myeloid (monocyte/macrophage) cells.

**Supplementary Figure 5: Monitoring weekly blood HIV titers in infected, dually engrafted mice.** Graphs show (y-axis) qPCR-based HIV copy number in weekly sampled cheek blood up to 4 weeks post-infection. (left) infected, (right) uninfected animals.

## Materials and Methods

### Construction and maintenance of fluorescent reporter iPSC

The iPSC lines were constructed at the Black Family Stem Cell Institute, using the well characterized WTC11 line (41) as starting point. The Cre recombinase-dependent dual fluorescent MSE2104 iPSC line was generated by CRISPR modification of the WTC11 line to insert a dsRed-to-eGFP cassette into the AAVS1 locus within the PPP1R12C gene (chromosomal location 19q13.4-qter) under the control of a CAG promoter. The MSE2104 iPSC line was maintained by culturing in feeder-free condition incomplete mTeSR E8 medium (StemCell Technologies) in a humidified incubator (5% CO2, 37°C) with medium changed every 1-2 days. Cells were passaged approximately every seven days dissociating the cells with 0.5mM EDTA in DPBS and plated onto 6-well plates (Corning) coated with growth factor-reduced Matrigel (1mg/mL; BD Biosciences) in mTeSR E8 medium supplemented with ROCK inhibitor Thiazovivin (Tocris). Media was switched to mTeSR E8 only medium the next day.

### iPSC differentiation into hematopoietic progenitor cells and microglia in vitro

HPCs and iMGs were differentiated according to published protocol (43–45, 50). On day 0, iPSCs were passaged in mTeSR-E8 medium for a density of 20-40 colonies of 100 cells each per 35mm well. On day 1, the cell medium was switched to STEMdiff Hematopoietic Kit (StemCell Technologies) Medium A. On day 4, the cell medium was switched to Medium B, in which the cells were differentiated into HPCs for 6-8 days. HPCs began to differentiate from the periphery of flattened endothelial cell colonies. Fully differentiated HPCs began to detach from the colonies to become suspended in medium B. On days 10-12, HPCs were harvested by collecting the medium and cells using a serological pipette. HPCs were validated by staining them with 1:100 dilution of CD34 (anti-human FITC, Biolegend), CD43 (anti-human PE, Biolegend), and CD45 (anti-human APC, BD Biosciences). Harvested HPCs were used for further differentiation in vitro or for HPC transplantation into neonatal mice (see below). The in vitro differentiation took 28-30 days in an iMG medium consisting of DMEM/F12, 2X insulin-transferrin-selenite, 2X B27, 0.5X N2, 1X glutamax, 1X non-essential amino acids, 400 mM monothioglycerol, and 5 mg/mL human insulin that was freshly supplemented with 100ng/mL IL-34, 50ng/mL TGFb1, and 25 ng/mL M-CSF (Peprotech). Differentiated iMGs were validated by measuring P2YR12 (anti-human PE dazzle 594, Biolegend), CD11b (anti-human APC, Biolegend), CXCR3 (anti-human BV650, Biolegend), CD45 (anti-human Alexa Fluor 700, Biolegend), CD206 (anti-human BV785, Biolegend), and CD14 (anti-human APC Fire 750, Biolegend).

### HIV-1 JRFL Cre clone construction and cell-free HIV-1 virion production

The HIV-1 JRFL clone is a full-length molecular clone of HIV-1 based on NL4-3 (57, 88) that expresses the JRFL envelope. Cre is inserted in place of the *nef* gene, and Nef expression is restored by a downstream internal ribosome entry site (IRES) (81). Plasmid was amplified in Stbl2 electrocompetent *E. coli* (NEB) and isolated using Qiagen Midi-kit. The human epithelial 293T cell line was used to produce HIV-1 virions. 293T cells were maintained in Dulbecco’s modified Eagle medium (DMEM; Sigma) containing 10% heat-inactivated fetal bovine serum (Sigma), 100 U/ml of penicillin (Gibco), 10 U/ml of streptomycin (Gibco), and two mM glutamine (Gibco) (complete DMEM). Cell-free virus particles were produced by transfection of 293T cells in a 10 cm dish using polyjet (Signajen) per the manufacturer’s protocol. Virus supernatant was harvested 48h post-transfection and filtered with 0.45 μm filter, concentrated by high-speed centrifugation (Sorvall ST 40R centrifuge; Thermo Fisher Scientific) at 100,000g for two h at 4°C. The pelleted virus was resuspended in DPBS, aliquoted, and stored at −80°C.

### HIV-1 p24 ELISA assay

Viral stocks were quantified by NCI HIV-1 p24 ELISA kit. Corning 96-well flat-bottomed plates were coated with anti-p24 capture antibody in 0.1 M NaHCO_3_ overnight at 4°C. Plate was blocked with 1% nonfat dry milk (Lab Scientific) for one h. The plate was then loaded with p24 standard titrations and experimental virus supernatant treated with 1% Empigen. The dish was incubated for two h at room temperature or overnight at 4°C and then washed six times with 1x TBS-0.05% Tween (TBST). Alkaline phosphatase-conjugated mouse anti-HIV p24 (Cliniqa) was added (1:8000 in TBST 20% sheep serum) and incubated for 1h, followed by 6 TBST washes with TBST. The plate was developed with Sapphire Substrate (Tropix), and luminescence was quantitated on a FluoStar Optima plate reader. HIV p24 level was calculated using Prism software (GraphPad), using nonlinear standard curve regression.

### iPSC-microglia and hu-PBMC xenograft mouse model

All procedures were performed in strict accordance with IACUC protocol at the Icahn School of Medicine at Mount Sinai. Neonatal immunocompromised mice (C;129S4-Rag2tm1.1Flv Csf1tm1(CSF1)Flv Il2rgtm1.1Flv/J, JAX ID# 014593) at age P0-P2 were taken out of their home cage and placed on sterile surgical drape overlying a cooling block for 2-3 min to induce hypothermic anesthesia. ICV injection of HPC was performed using a 30G needle fixed to a 10 μl Hamilton syringe. Each mouse received 400-500K HPCs at four cranial surface coordinates at two different depths, totaling eight different sites (44). The HPCs were resuspended in 1x DPBS at 50-62.5K cells/μl for injection. Injected mice were allowed to recover on heating pads covered with sterile surgical drapes before being returned to their home cages. Mice were weaned from their mother at P21.

At 6-10 weeks, mice hosting the central xenograft were intraperitoneally injected with human PBMCs. PBMCs were obtained from deidentified HIV-1 negative healthy blood donors (New York Blood Center), purified by Ficoll (HyClone) density gradient centrifugation, and maintained in RPMI 1640 medium (Sigma) containing 10% heat-inactivated fetal bovine serum (Sigma), 100 U/ml of penicillin (Gibco), 10 U/ml of streptomycin (Gibco), and two mM glutamine (Gibco) (complete RPMI). To minimize the donor variability effect, we used the same PBMC donors to inject all nine mice used for this study. PBMCs were activated with phytohaemagglutinin-L (PHA-L; 2 μg/ml, Sigma) and IL-2 (50 IU/ml, Roche) for three days co-cultured with irradiated feeder PBMCs. Cells were harvested, counted, and 10^7^ cells resuspended in 200 μl 1x PBS were intraperitoneally injected into each mouse. One week after PBMC injection, engraftment was measured weekly by quantifying human CD45+ cells in each mouse’s peripheral blood through fluorescence-activated cell sorting (FACS) on Attune flow cytometer (ThermoFisher). On average, mice have successfully engrafted with PBMC ∼4 weeks after the initial injection.

### PBMC engraftment FACS analysis

Attune flow cytometer (ThermoFisher) was used to measure the level of human PBMC engraftment in our iPSC-microglia and hu-PBMC xenografted mice. The cellular layer separated from the plasma of peripheral blood was treated with ACK lysis buffer (Gibco) to remove red blood cells. Isolated white blood cells were stained with LIVE/DEAD fixable stain (Invitrogen) at a concentration of 1:1000 in FACS buffer (2 mM EDTA, 2% FBS in DPBS) to detect live cells. Cells were incubated for 30min at 4°C and then washed with FACS buffer. Cells were stained with 1:100 concentration of CD45 (anti-human PE-Cy7, Biolegend), CD45 (anti-mouse Pacific Blue, Biolegend), CD3 (anti-human APC eFluor780, eBiosciences), CD4 (anti-human APC, Biolegend), and CD8 (anti-human PerCP-Cy5.5, Biolegend) for 30min at 4°C. Stained cells were washed with FACS buffer and fixed in 4% (w/v) PFA for FACS analysis.

### HIV-1 infection of iPSC-microglia and hu-PBMC xenografted mice

For HIV-1 infection, each mouse was injected with 250ng HIV-1 p24 antigen either intracranially or intraperitoneally. For ICV infection, a rodent stereotaxic rig mounted with a micro pump (Stoelting) and Hamilton syringe fitted with a 30G needle was used to inject HIV-1 bilaterally into the PFC (1 μl per hemisphere) (91). The coordinates for injection were as follows: +1.5 mm anterior/posterior, ±0.5 mm medial/lateral, and 1.5 mm dorsal/ventral. The virus was injected per hemisphere at 0.25 μl min−1, and four additional minutes were allowed before syringe removal. Mice peripheral blood was collected and analyzed weekly for evidence of peripheral infection. Mice were sacrificed four weeks post-infection, and tissue samples were harvested and analyzed.

### Peripheral HIV-1 infection qPCR analysis

Peripheral blood collected from mice was centrifuged at 10,000g for 10min to separate plasma from cells. RNA was isolated from plasma using QIAamp Viral RNA Mini kit (Qiagen) and quantified using NanoDrop Spectrophotometer (ThermoFisher). RNA was reverse transcribed to cDNA using the High-Capacity RNA-to-cDNA kit (ThermoFisher). A Custom TaqMan Gene Expression RT-PCR assay designed for the *gag-pol* region (Assay ID: AP7DXHY; ThermoFisher) was then used on the cDNA to quantify the HIV-1 viral copy number. A series of 10-fold dilutions of measured HIV-1 target RNA fragments derived from the HIV-1 NL4-3 clone was included in each assay to generate a standard curve to derive the HIV-1 copy number.

### Immunohistochemistry of mice brain sections and Confocal Microscopy

Mice were anesthetized with isofluorane and monitored for loss of consciousness. Mice that did not respond to toe pinch were cervically dislocated, and their brains dissected. Brains were drop fixed in 4% (w/v) PFA for 24h. Fixed brains were cryoprotected in 30% (w/v) sucrose until they sank to the bottom of the solution for at least 48h. Brains were cut coronally or sagittally at 20-40 μm thickness using a sliding microtome cooled with dry ice. Free-floating tissue sections were collected in 1x DPBS and 0.05% sodium azide. For immunohistochemistry staining, tissues were blocked in 1x DPBS, 0.1% Triton X-100, and 1% BSA for one h at room temperature. Tissues were incubated in primary antibodies diluted in 1x DPBS and 1% BSA overnight on a shaker at 4°C. Tissue sections were washed with DPBS x 3 at room temperature and incubated in fluorophore-conjugated secondary antibodies either for one h at room temperature or overnight at 4°C. Tissues were washed with DPBS x 3 at room temperature and then stained with DAPI (Sigma Aldrich). Tissues were washed with DPBS and then mounted on charged glass slides. Immunofluorescent sections were visualized and imaged using either Zeiss LSM780 or Zeiss LSM980 with airyscan2 confocal microscopes. Brightness and contrast settings were slightly adjusted for better visualization of some images. Primary antibodies: mouse anti-human nuclei (Ku80 1:100; Abcam, ab79220), mouse anti-human nuclei (HuNu 1:50; Millipore, mab1280), rabbit anti-Iba1 (1:100; Wako, 019-19741), goat anti-Iba1 (1:100; Abcam ab5076), rabbit anti-P2ry12 (1:500; Sigma; HPA014518).

### RNAScope Fluorescence in situ Hybridization

Fixed and cryoprotected mice brain tissue sections were cut coronally or sagittally at 10-20 μm thickness using a freezing sliding microtome. Tissue sections were mounted on charged slides and dried for 10 min at 60°C. The slides were processed per the RNAscope Multiplex Fluorescent v2 protocol (Advanced Cell Diagnostics). Mounted tissue sections were incubated with an RNA-specific probe against the HIV-1 *gag-pol* gene (Probe-V-HIV1-gagpol-sense-C2, Advanced Cell Diagnostics) overnight at 40°C. Amplifier sequences were polymerized to the probes and treated with Opal 570 dye. Imaging was performed on Zeiss LSM780 confocal microscope.

